# All-or-none disconnection of pyramidal inputs onto parvalbumin-positive interneurons gates ocular dominance plasticity

**DOI:** 10.1101/2020.12.30.424871

**Authors:** Daniel Severin, Su Z. Hong, Seung-Eon Roh, Shiyong Huang, Jiechao Zhou, Michelle C. D. Bridi, Ingie Hong, Sachiko Murase, Sarah Robertson, Rebecca Haberman, Richard Huganir, Michela Gallagher, Elizabeth M. Quinlan, Paul Worley, Alfredo Kirkwood

## Abstract

Disinhibition is an obligatory initial step in the remodeling of cortical circuits by sensory experience. Our investigation on disinhibitory mechanisms in the classical model of ocular dominance plasticity uncovered an unexpected novel form of experience-dependent circuit plasticity. In layer 2/3 of mouse visual cortex monocular deprivation triggers a complete, “all-or-none”, elimination of connections from pyramidal cells onto nearby parvalbumin-positive interneurons (Pyr➔PV). This circuit plasticity is unique as it is transient, local and discrete. It lasts only one day, and it does not manifest as widespread changes in synaptic strength, rather, only about half of local connections are lost and the remaining ones are not affected in strength. Mechanistically, the deprivation-induced loss of Pyr➔PV is contingent on a reduction of the protein neuropentraxin2 (NPTX2). Functionally, the loss of Pyr➔PV is absolutely necessary for ODP. We surmise, therefore, that this “all-or-none” loss of local Pyr➔PV circuitry gates experience-dependent cortical plasticity.

## INTRODUCTION

Experience during a postnatal critical period is essential to properly shape the functional connectivity of cortical circuits. A canonical model of cortical plasticity is the shift in ocular dominance following monocular deprivation (MD), which biases responses towards the nondeprived eye. Prior research established that MD-induced changes result from the reorganization of excitatory glutamatergic synapses onto excitatory pyramidal neurons, which is in turn regulated by an inhibitory GABAergic network composed of parvalbumin-positive inhibitory interneurons (PVs). The current consensus is that a reduced, permissive, level of inhibition from PV circuits in cortical layer 2/3 is required for plasticity at downstream excitatory synapses, and that inhibition above or below the permissive range constrains the response to MD (Feldman, 2000; Hensch and Quinlan, 2018; Jiang et al., 2005). Although the notion that rapid cortical disinhibition precedes and initiates plasticity of glutamatergic connectivity is well established (Barnes et al., 2015; Kuhlman et al., 2013), and decades old (Hendry and Jones, 1986; Hickmott and Merzenich, 2002; Levy et al., 2002), the underlying cellular mechanisms remain unclear.

Disinhibition of excitatory cortical neurons could be achieved indirectly, for example by suppressing PV activity via enhancing inhibition from other interneurons through cholinergic neuromodulation (Froemke et al., 2007; Letzkus et al., 2011); but more directly, and likely more effectively, by reducing the excitatory input onto PVs (Gu et al., 2013; Gu et al., 2016; Kuhlman et al., 2013; Sun et al., 2016). Our current investigation uncovered a unique novel form of experience-dependent plasticity that regulates the connectivity between pyramidal neurons and PVs. We found that the initial response to monocular deprivation is the functional and structural elimination of ~50% of these connections. In contrast to the outcome of known mechanisms of synaptic plasticity that manifest in widespread graded changes in synaptic strength, the loss of pyramidal-PV connectivity occurs in a discrete, “all-or-none”, fashion: whereas a subset of connections become completely eliminated, the persistent connections have normal strength. This disconnection is not only rapid, but it is transient, affects only very local pyramidal-PV pairs, and, importantly, manipulations that promote/prevent this disconnection also promote/prevent shifts in ocular dominance. We surmise, therefore that the rapid and transient disconnection of discrete subsets of PV circuits enables the subsequent Hebbian and homeostatic modification of glutamatergic circuitry.

## RESULTS

### Brief MD transiently reduces local connectivity between Pyrs→PVs

We evaluated how monocular deprivation (MD) affects the connectivity between pyramidal neurons (Pyr) and PV interneurons (Pyrs→PVs) in mouse visual cortex (V1) layer 2/3. To that end, we performed visualized whole cell-paired recordings in slices containing the monocular zone of V1 of mice expressing Td tomato in PVs (Fig. 1A,B. See methods). Consistent with previous studies (Gu et al., 2013; Lu et al., 2014), in juvenile mice (p24-p30) the connection probability in proximate Pyr→PV pairs (less that 50 μm apart) was high (p~0.5) in slices from normal reared (NR) mice. In mice subjected to MD for 1 day (MD1) connection probability was reduced in the V1 contralateral to the deprived eye (Fig. 1C), but only for the pairs less that ~25 μm apart. A logistic regression analysis confirmed the significance of these differences (|Z|=3.228; p=0.0012), including the interaction between experience and Pyr-PV distance (|Z|=2.435; p=0.0149). In contrast, MD1 did not affect the connectivity in the nondeprived (ND) cortex ipsilateral to the deprived eye (NR-ND: p=0.6005). Using this ND cortex as a control we found the disconnection in deprived cortex is transient: the differences between cortices are significant after 1 day, but not after 2 and 3 days of MD (Logistic regression: (|Z|=2.370; p=0.0178; interaction between experience and days |Z|=2.020; p=0.0434; Fig. 1d). Importantly, the amplitude of the unitary excitatory postsynaptic current (uEPSC) of the proximal pairs that remained connected (within 12-24 microns) was not affected by MD (Fig. 1E,F). Moreover, MD did not affect crucial determinants of the uEPSC amplitude, including the quantal size, failure rate, and the size and replenishment rate of the readily releasable vesicle pool (Supplementary Fig. 1). Finally, we found that in the binocular zone MD(1) also reduces the Pyr→PV connection probability, from 0.69±0/.09 (N=5 mice, 29 pairs) in NR mice to 0.40±0.09 in MD mice (N=5,42; p=0.029. Data not shown).

**Fig. 1.**
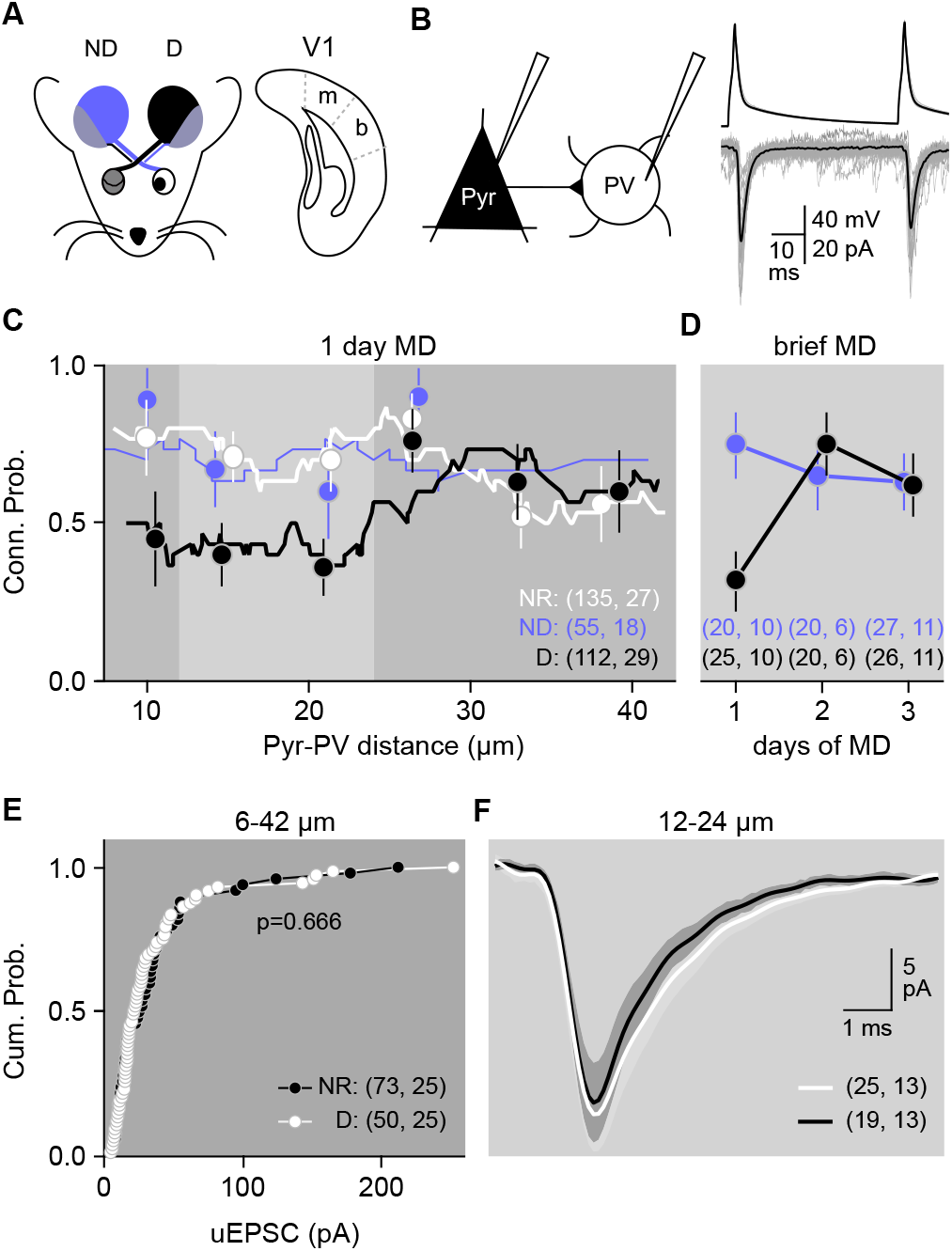
MD transiently eliminates functional connections between local Pyr→PVs. **A,** Paired whole cell recordings from cortical slices containing deprived (D), nondeprived (ND) or normal-reared (NR) monocular V1. **B,** Example of uEPSCs (individuals: gray; average: black) evoked in a PV-IN (PV) by action potentials produced in pyramidal neuron (Pyr). **C**, probability that Pyr→PV pairs are synaptically connected versus distance between somas. Lines: 21 points running averages; symbols: binned averages over 10 μm (NR: white; D(1 day): black; ND: blue). **D**, Transient changes in functional connectivity during MD. The difference in pair connectivity between the D and ND hemispheres was significant following 1 day of MD only (χ^2^[3×2]=10.84 p=0.0547; F-test MD(1) p=0.0139; F-test MD(2) p=0.7311; F-test MD(3) p>0.9999). **E,** Comparable distributions of the uEPSC magnitude of connected pairs in NR (white circles, and D (black circles). KS-test: p=0.666. **F**, MD does not affect the uEPSC magnitude of the connected pairs separated by > 12 μm and < 24 μm (light grey zone in c). NR: 28.0 ±3.5 pA; D(1): 24.6 ±3.1 pA; MW-test: U=222.5, p=0.7294). Traces are averaged uEPSCs (± 95% C.I) for all connected pairs. The number of pairs and mice is indicated in parenthesis in c,d,f.

Layer 2/3 PVs are also driven by excitatory inputs from layer 4 (Xu and Callaway, 2009). Since the probability of detecting connected intralaminar Pyr→PV pairs is typically very low (2 out of 20; not shown), we examined the effects of MD by computing the amplitude of maximal compound EPSCs as a measure of total strength in NR and MD mice (Gu et al., 2013; Morales et al., 2002). MD(1) reduced the amplitude of the maximal EPSC recruited electrically from layer 2/3 but not the EPSC recruited from layer 4 (Fig. 2c-g). Next, we evaluated changes in the ratio of EPSCs evoked in nearby Pyr and PVs by optogenetic stimulation of layer 4 inputs. MD(1) did not affect the EPSC ratio in PV/Pyr pairs (Fig. 2A,B). Because one day of MD does not affect excitatory inputs onto layer 2/3 pyramidal cells (Goel and Lee, 2007; Kuhlman et al., 2013), the absence of changes in the argues against alterations in layer 4 inputs to layer 2/3 PVs. Finally, we evaluated MD-induced changes in the outputs of PVs: PV➔PV and PV➔Pyr. MD(1) did not affect the amplitude of the maximal IPSC recorded in PVs (Supplementary Fig. 2A-C), the proportion of connected PV➔Pyr pairs, nor the amplitude of the unitary IPSCs in Pyrs (Supplementary Fig. 2D-G). In sum, a primary effect of brief MD on Layer 2/3 PV circuitry is the selective and complete, yet transient, “all or none” elimination of approximately half the functional connections made by nearby pyramidal cells, without affecting the potency of the remaining connections.

**Fig. 2.**
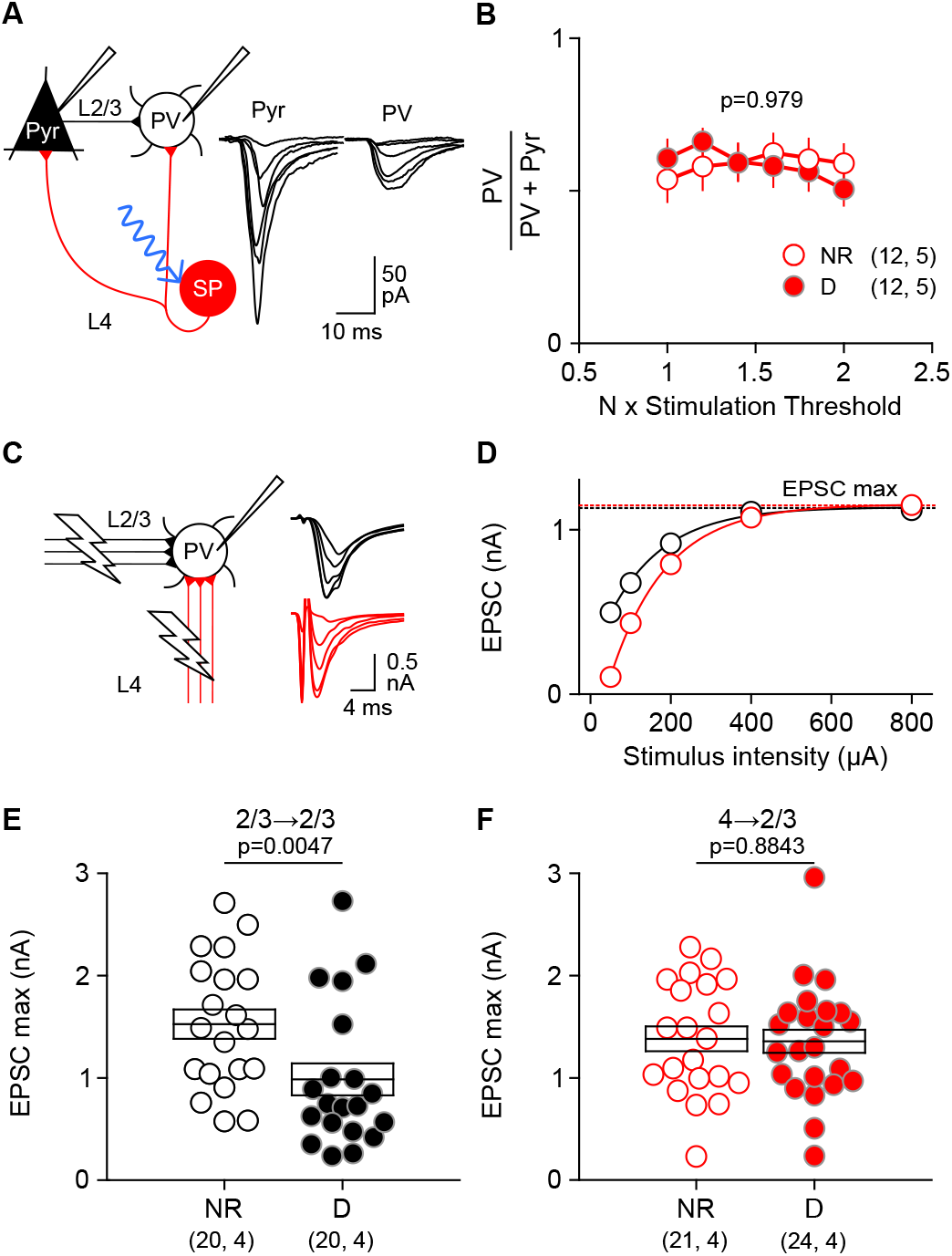
MD(1) selectively reduces intralaminar excitatory inputs onto layer 2/3 PVs without affect ascending inputs from layer 4. **A,** Example EPSCs evoked in neighboring PVs and Pyrs by activating layer 4 inputs with optogenetic stimulation of increasing intensities (see methods for details), The response ratio (PV/PV+Pyr) was similar across stimulation intensities, and similar in pairs from NR (open circles) and D (filled circles) mice (ANOVA F[1,128]=0.0007, p=0.9794; interaction F[5,128]=0.49, p=0.7844). Averages are shown as circles ±. s.e.m. **C**-**D**, Maximal EPSCs (EPSCmax) evoked by electrical stimulation of increasing intensity applied to layer 2/3 (black) or layer 4 (red), shown in **C**, were computed from input/output plots (shown in **D**). **E,** MD(1) reduced the EPSCmax evoked from layer 2/3 (NR:1.525±0.143 nA, D: 0.986±0.156 nA; MW-test U=97, p=0.0047). **F**, MD(1) did not reduce the EPSCmax evoked from layer 4 (NR: 1.380±0.122 nA; D:1.356 ±0.113 nA; t-test t=0.1464, p=0.8843. **G**, MD(1) reduces the ratio of EPSCmax evoked from layer 2/3/ layer 4. Circles in **E,F,G**: EPSCmax from individual cells; boxes: averages ± s.e.m. The number of cells or cell pairs and mice is indicated in parenthesis in **B, E,F,G.**

### Evidence that MD induces a transient structural loss of Pyr→PV connections

The “all or none” elimination of functional Pyr→PV connectivity is reminiscent of synaptic pruning. Hence it suggests a loss of synaptic structure rather than a reduction in synaptic strength. To test this idea, we first considered the possibility that MD might “silence” Pyr→PV connections by inducing the removal of synaptic AMPA receptors, as has been documented in pyramidal neurons. To that end we compared the AMPA/NMDAR EPSC ratio, a crude indicator of changes in silent synapses, in PVs from deprived and non-deprived cortices. We detected no differences following MD1, arguing against a silent synapse scenario (Fig. 3A,). Next, we evaluated changes in the number excitatory synapses onto PVs using immunocytochemistry to visualize the vesicular glutamate transporter 1 (VG1) as a marker for intracortical glutamatergic terminals. After 1 day of MD, the average number of VG1 puncta in the soma and proximal dendrites of PVs was reduced in deprived relative to non-deprived monocular zone, but there was no difference after 3 days of MD (Fig. 3B,C). These results suggest that MD causes a transient structural elimination of excitatory synapses onto PVs.

**Fig. 3.**
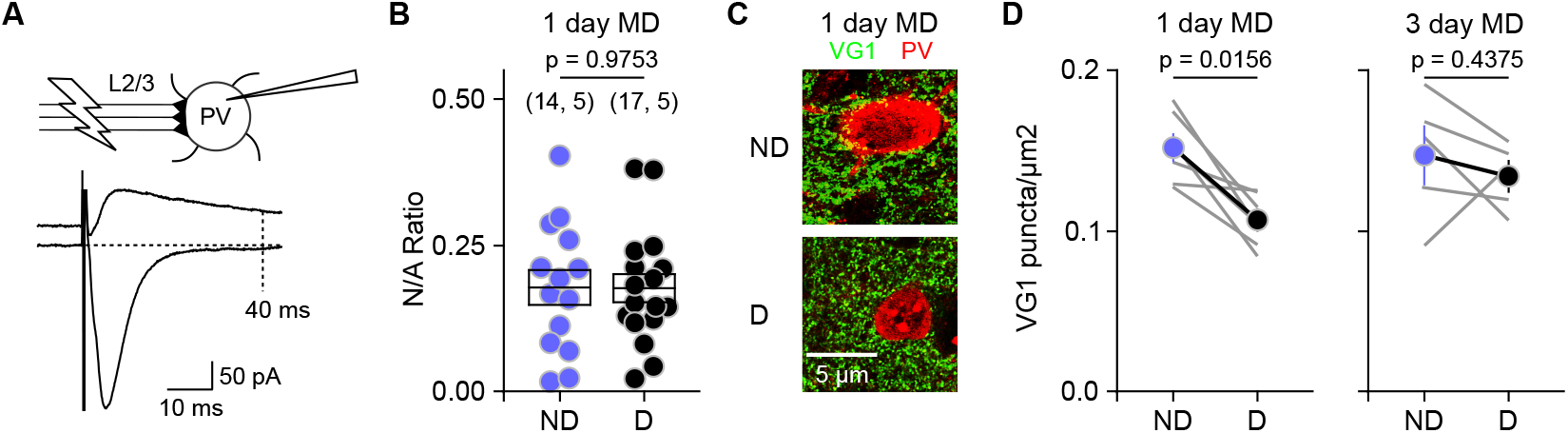
MD(1) induces a transient loss of structural Pyr→PV connections. **A-B,** Evidence against change in silent synapses. **A**, Schematic for measurement of NMDAR and AMPAR epscs. **B**, MD(1) does not affect the NMDAR/AMPAR (N/A) EPSC amplitude ratio (t-test F[29]=0.03165, p=0.9750). The number of cells and mice is indicated in parenthesis. **C-D,** Immunohistochemical analysis of the MD-induced changes in the VGlut1 puncta density on the soma and proximal dendrites of layer 2/3 PV-INs. **C**, Example cases (projection of 3 image planes) showing PV in red, VG1 in green. Colocalized puncta are depicted in yellow. **D**, Quantification of colocalized PV+VG1. The grey lines connect results of the deprived (D) and non-deprived (ND) monocular V1 of the same mouse after MD for 1 day (left) or 3 days (right). Averages are shown as circles connected by black lines ± s.e.m.

### NPTX2 but not NRG1 mediates the disconnection of Pyrs→PVs by MD

Attractive candidate mechanisms to mediate the disconnection of Pyr→PV synapses by MD include neuregulin1 (NRG1) and neuropentraxin2 (NPTX2) signaling. Both molecules are secreted by pyramidal neurons in an activity-dependent manner and promote/stabilize AMPARs at excitatory synapses onto PVs in multiple brain regions, including the visual cortex (Fazzari et al., 2010; Gu et al., 2013; Gu et al., 2016; Tsui et al., 1996). A reduced secretion of NRG1 during MD has been associated with diminished excitatory input onto PV-INs (Sun et al., 2016). We therefore tested whether supplemental NRG1 (1μg subcutaneous (Gu et al., 2016)) prevents the disconnection of Pyr→PV during MD(1). Systemic NRG1 did not inhibit the decrease in Pyr→PV connection probability in deprived V1 after MD1 (Fig. 4A). Next, we examined how exogenous NRG1 affects excitatory inputs onto layer 2/3 PVs in slices from MD1 mice. To that end we recorded in the same PVs the EPSCs evoked by layer 2/3 and layer 4 stimulation. NRG1 application specifically enhanced the inputs from layer 4 (Fig. 4B), which are normally not affected by MD (see Fig. 2). A comparable enhancement of layer 4 inputs was also observed in normal reared V1b (Fig. 4 C). Together this argues against a role for NRG1 in the elimination of Pyr→PV connections in layer 2/3 by MD1.

**Fig. 4.**
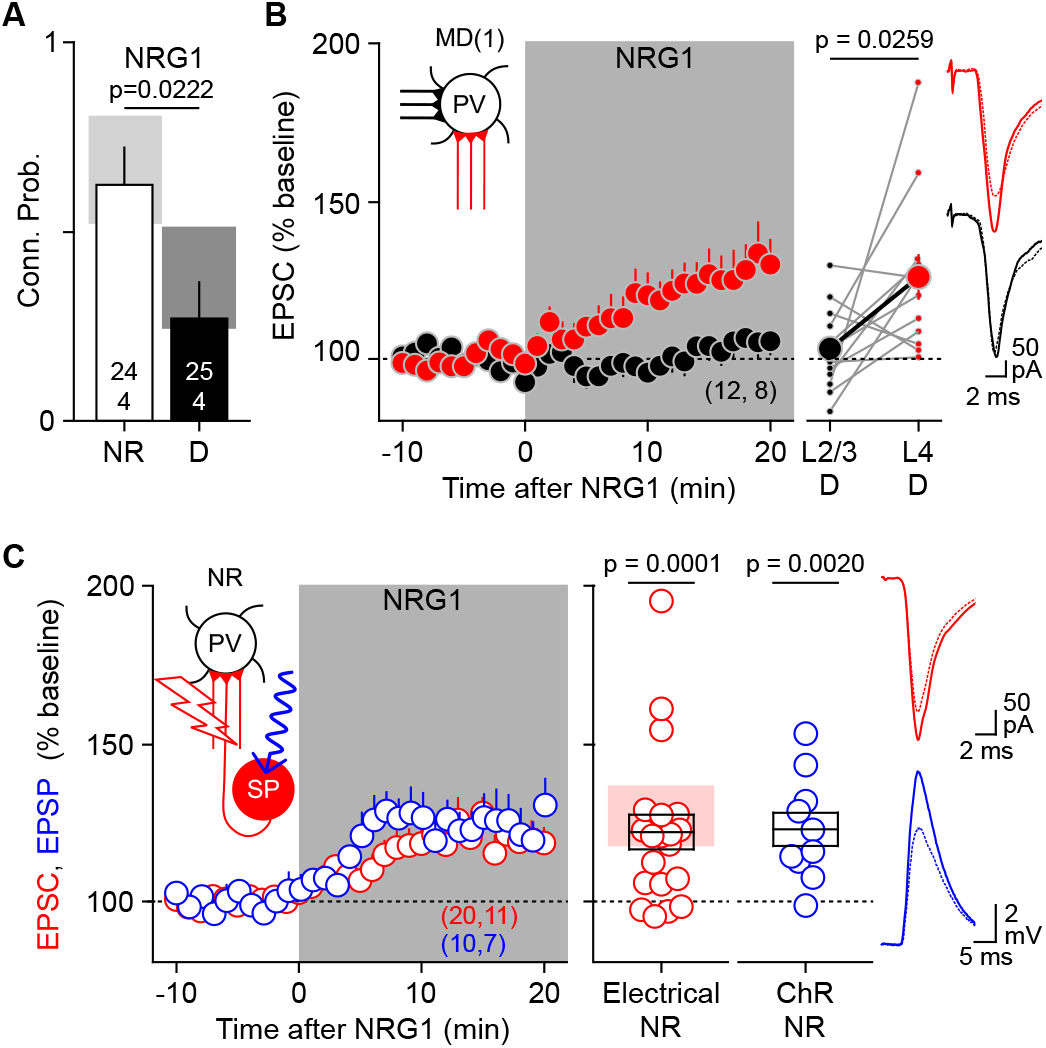
NRG1 does not affect inputs onto PVs from layer 2/3 Pyrs. **A,** In mice between p21-25 systemically injected with NRG1 (1μg/mouse subcutaneous, 1X prior to eye suture and 1X 1h before sacrifice; MD(1) reduces the Pyr→PV connection probability. Gray bars indicate the 95% CI in non-injected mice (data from Fig 1). **B**, In slices from MD mice, superfusion of NRG1 (5 nM grey zone) selectively potentiated EPSCs evoked from layer 4 (red symbols: 126.0±7.3% at 10-20 min. Wilcoxon test W=78, p=0.0005) without affecting EPSCs evoked from layer 3 (black symbols: 103.3±3.9%, Wilcoxon test W=16, p=0.5547). Left: average time course; right: grey lines connect EPSCs evoked by layer 4 and layer 3 stimulation in the same cell; the black lines connect averages ± s.e.m. **C,** NRG1 (5 nM grey zone) also potentiates EPSCs evoked by layer 4 stimulation in slices from normal-reared mice. Red symbols: electrically-evoked EPSCs; blue symbols: optogenetically-evoked EPSPs in mice expressing ChR2 in layer 4 (see methods). Left: average time course. Right: changes in individual cells; boxes: averages ± s.e.m; shaded red box: 95% C.I. for layer 4-evoked EPSCs in deprived slices (from 4B). Number of cells or cell pairs and mice is indicated in parenthesis in **A-C**.

Genetic ablation of NPTX2 (also known as NARP) has been shown to result in the all-or-none elimination of a substantial proportion of Pyr→PV connections (Gu et al, 2013). To examine the role of NPTX2 in the disconnection induced by MD, we monitored how MD changes the release NPTX2 *in vivo*. NPTX2 release was monitored by viral expression of NPTX2 fused with Super Ecliptic pHlurorin (SEP) which fluoresces only in the extracellular compartment (Martineau et al., 2017). In wild type mice virally transfected with AAV-CaMKII-NPTX2-SEP in V1, two-photon imaging in the superficial cortical layers revealed discrete puncta (2-3 μm) likely representing extracellular aggregates of NPTX2-SEP (Fig. 5b). One day of MD substantially reduced the density and intensity of these puncta, which returned to normal levels by the 2^nd^ day of MD (Fig. 5-D). A reduction in NPTX2 protein content in V1 after one day of MD is also revealed by quantitative immunochemistry (supplementary Fig. 4). These results suggest that MD transiently reduces the synaptic expression of NPTX2.

**Fig. 5.**
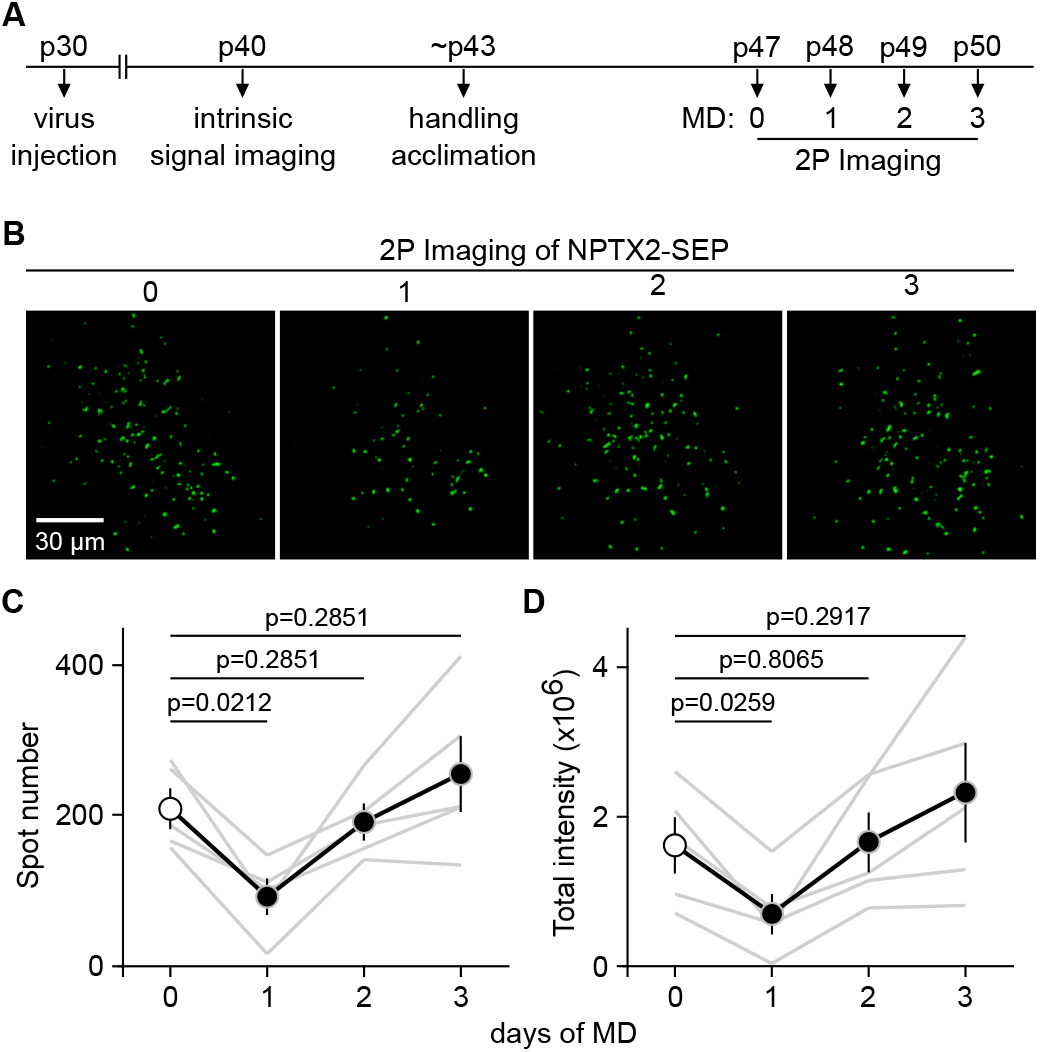
MD transiently reduces surface NPTX2 in L2/3 V1. **A**, Time schedule of the experiment. AAV-CaMKII-NPTX2-SEP was injected into the visual cortex of wild-type mice at P30. Intrinsic signal imaging was carried out at P40 to identify V1. Around P47, live animals were imaged in layer 2/3 with 2-photon microscopy. **B,** Example images from the same mouse before and after 1, 2, and 3 days of MD reveals a brief reduction of the number of NPTX2-SEP puncta at day 1. **C,** Quantification of NPTX2-SEP puncta number in L2/3 (80 −180μm) indicates a significant and transient reduction at day 1 (Repeated measures ANOVA F[1.508, 6.031]= 12.61, p=0.0087). **D,** The quantified total fluorescence intensity (in arbitrary units) of the NPTX2-SEP puncta is transiently reduced by 1 day MD (Repeated measures ANOVA F[1.132, 4.526]= 7.209, p=0.0466. Comparisons indicated in **C,D** used the Holm-Sidak’s multiple comparison test. Grey lines in **C,D** represent data from individual mice; N = 5 mice; circles and black lines, average ± s.e.m.

To test whether an MD-induced reduction in NPTX2 content is necessary for the Pyr➔PV disconnection, we exploited the fact that the effects of MD on Pyr➔PV-IN inputs have a critical period. The loss of connectivity induced by one day of MD are robust up to postnatal day 50 (p50) but cannot be elicited at ~p100 (Fig. 6A). We therefore tested whether overexpression of NPTX2 prevents the MD-induced Pyr➔PV disconnection in juveniles, and whether reducing the functionality of NPTX2 enables the disconnection in adults. In the first case we transfected V1 neonatally with AAV-CaMKII-NPTX2-SEP, in the second case we transfected V1 in adults with AAV2/2.CAMKII.dnNPTX2.EGFP to express a dominant negative form of NPTX2 that disrupts biosynthesis of the native NPTX1/2/R complex (Charbonnier-Beaupel et al., 2015). In juveniles (p21-p25) overexpressing NPTX2-SEP, the Pyr➔PV connectivity remained high after MD(1), comparable to age-matched normal-reared controls (Fig. 6B). As a control we confirmed reduced connectivity after MD(1) in juveniles transfected with the same serotype AA virus expressing GFP (AAV2-CaMKII-GFP). On the other hand, in post-critical period adults (~p110) the expression of dnNPTX2, which by itself did not affect the Pyr➔PV connectivity, enabled a substantial and significant disconnection of Pyr➔PV inputs following MD(1) (Fig. 6C). As in the case of critical period mice the amplitude of the remaining EPSCs was normal (Fig. 6C). These results support a model in which a reduction of NPTX2 is a necessary permissive factor for the MD-induced disconnection of Pyr→PV inputs in layer 2/3.

**Fig. 6.**
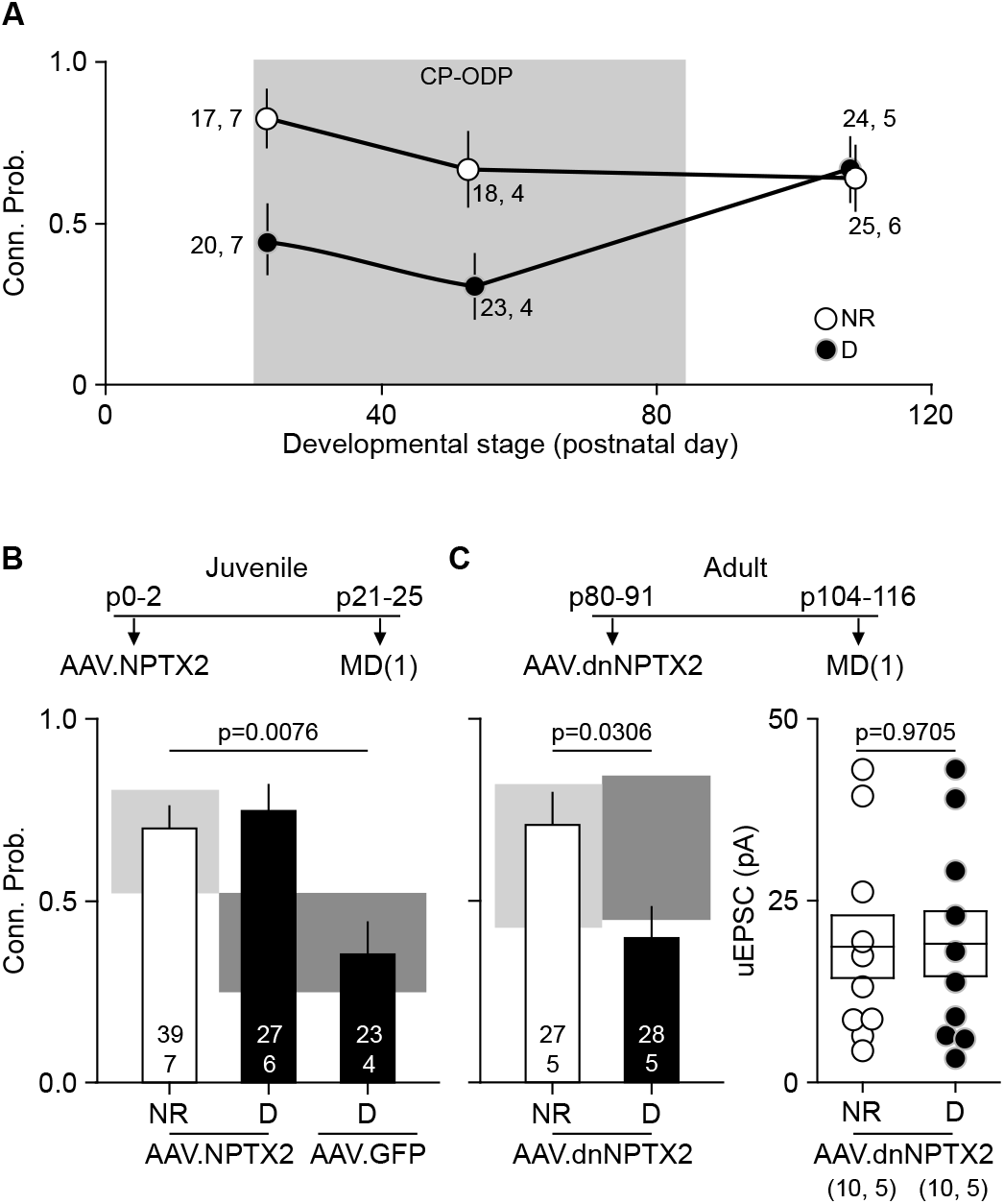
Manipulations of NPTX2 prevent/enable MD-induced Pyr→PV disconnection. **A**, Critical period for the MD-induced plasticity of Pyr→PV inputs. In mice younger than ~p60 MD but not in ~p110 mice, MD for 1 day reduces the EPSC max amplitude (red circles. 2-way ANOVA F[1, 106]=18.66, p<0.0001; interaction p=0.006) and the probability of connection (black circles. Logistic regression F[1,23]=8.835, p=0.0036; interaction F[1,23]=3.918, p=0.0500). Data shown as averages ± s.e.m (Open circles: NR mice; filled circles: D mice). **B**, Transfection with AAV2-CaMKII-NPTX2-SEP at ~ birth prevents MD(1)-induced disconnection of Pyr→PV inputs in juveniles. Top: experimental schedule. Bottom: Pyr➔PV connection probability in normal reared (NR), deprived (D) mice transfected with NPTX2-SEP, and deprived mice transfected with control virus (AA2-GFP). **C**, transfection of V1 with AAV2-dnNPTX2 in adult mice enables the MD(1)-induced disconnection of Pyr→PV inputs. Top: experimental schedule. Bottom: the connection probability in normal reared (NR) and deprived (D) transfected mice. For comparison, boxes in **B,C** depict the 95% CI range non-transfected mice (light grey: normal-reared; dark grey: deprived). The number of cells or cell pairs and mice is indicated in parenthesis in **A-C.**

### Manipulation of NPTX2 prevent/enables ocular dominance plasticity

Finally, we examined whether the manipulations of NPTX2 that prevent/enable the Pyr➔PV disconnections also prevent/enable the plasticity of ocular dominance (ODP) induced by MD. In mice, ODP induced by MD has a critical period with a time course parallel to the time course for MD-induced disconnections of Pyr➔PV inputs in layer 2/3 (Fig. 6A). ODP in the critical period is manifest as a rapid reduction of cortical responsiveness to the deprived eye in juveniles (younger than ~p35), and a delayed increase responsiveness to the non-deprived eye in young adults (younger than ~p70; (Cooke and Bear, 2014; Lehmann and Lowel, 2008). First, we asked how overexpression of NPTX2 following neo-natal AAV transfection (as in Fig. 6B), affects ocular dominance shift induced by brief MD in juveniles (3 days Fig. 7A). We used intrinsic signal imaging to evaluate cortical responses in the binocular zone of V1 as described (Hong et al., 2020; Kaneko et al., 2008). Cortical responses in mice are normally biased toward the contralateral eye and this bias is reduced by MD, reflected as a shift of the ocular dominance index (ODI, see methods). In NPTX2-transfected mice, the ODI for normal-reared and MD individuals was similar (Fig. 7B) and within the range for normal reared non transfected mice (Fig. 7, light gray bar). On the other hand, mice transfected with control AAV and subjected to MD(3) exhibited a reduction in ODI, comparable to the decrease in ODI induced by MD3 in non-infected controls Fig. 7b, dark gray bar). In a complementary study we determined that transfecting adult mice (>p110) with AAV-dnNPTX2 to reduce NPTX2 function (as in Fig. 6C), enables a juvenile-like decrease in ODI in response to MD. In these experiments we imaged V1b before and after MD(3) (Fig. 8A) and confirmed that the reduction in ODI was juvenile-like; that is, it resulted from a reduction in the contralateral response, not from an increase in the ipsilateral response (Fig. 8B-E). Altogether, the results indicate that a loss of NPTX2-dependent Pyr➔PV-IN connectivity is an obligatory early step in ODP.

**Fig. 7.**
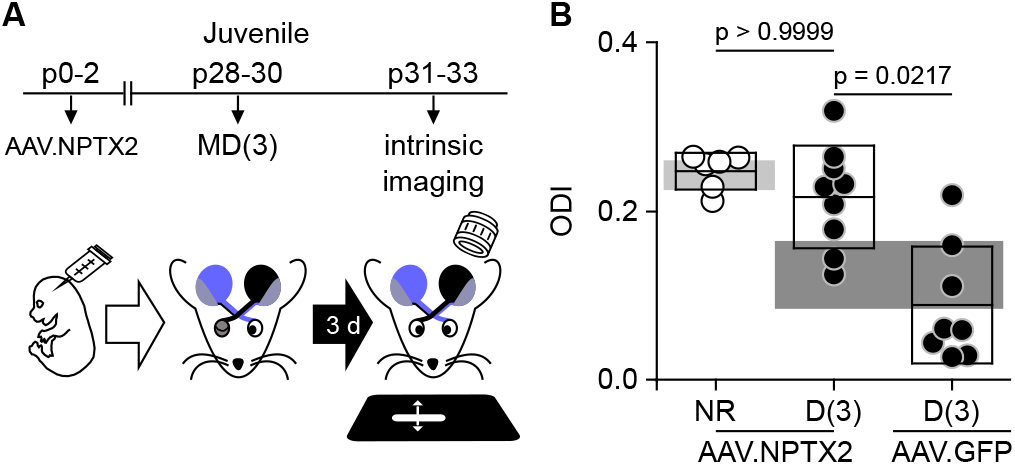
Overexpression of NPTX2 prevents ocular dominance plasticity in juveniles. **A,** Mice were transfected at birth (p0-2) with AAV2-CaMKII-NPTX2-SEP or AAV2-GPP. Following 3 days of monocular deprivation starting at p30-p31, intrinsic signal responses to stimulation in each eye were recorded in the V1 contralateral to the deprived eye (see methods). **b,** In the NPTX2 transfected mice (black circles), the ocular dominance index (ODI: see methods) after MD(3) was comparable to nondeprived transfected mice (white circles), and larger than in deprived mice transfected with control AAV2-GFP (black triangles). ANOVA F[3,23]=12.56, p=0.0019; NR_NPTX2_-D(3)_NPTX2_ p>0.9999; D(3)_NPTX2-D_(3)_GFP_ p=0.0217). For comparison, boxes in **b** depict the 95% CI range of ODI of 44 NR and 8 D non-transfected juvenile mice after MD(1) (light grey: normal-reared; dark grey: deprived).

**Fig. 8.**
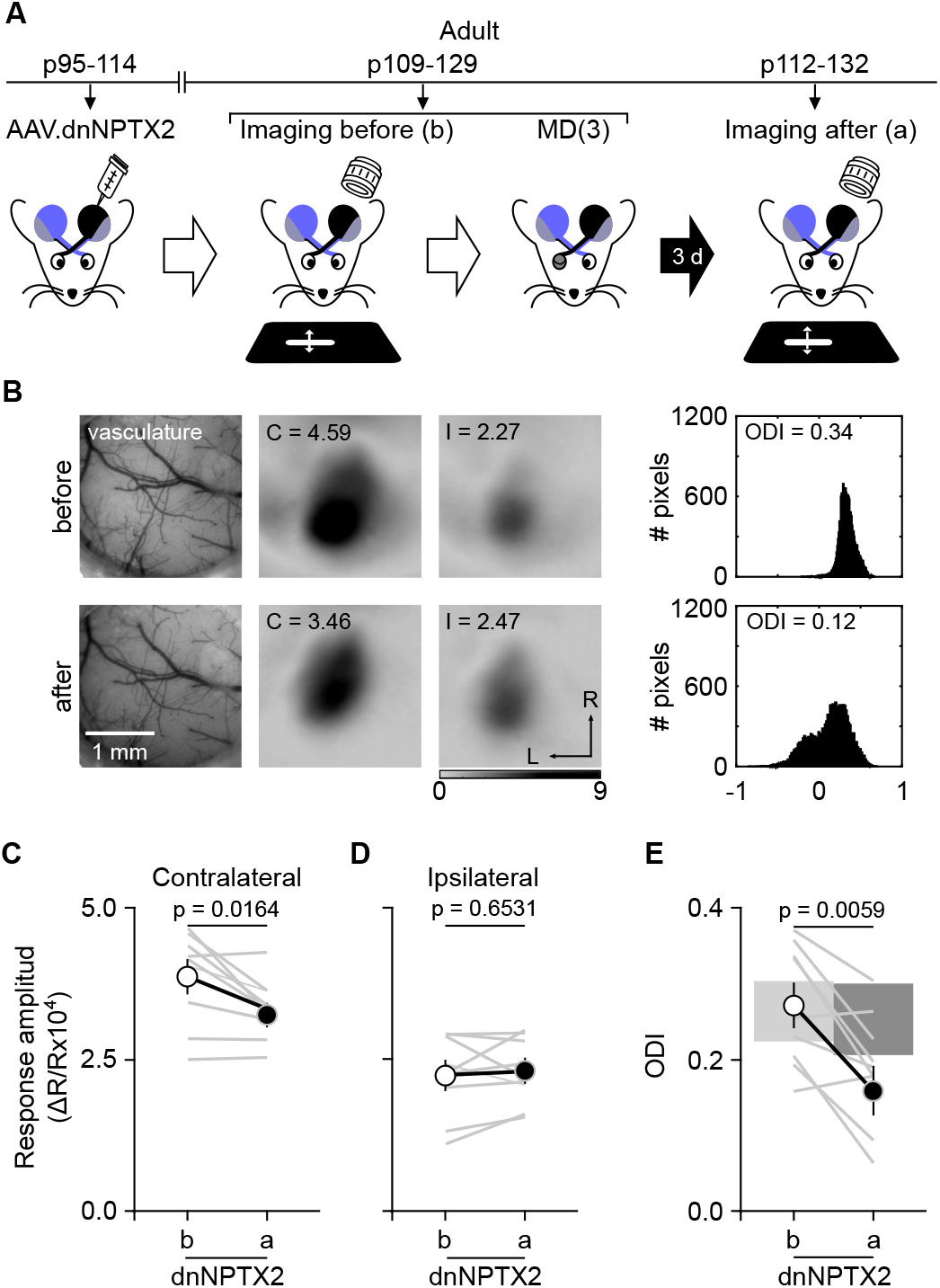
Expression of dnNPTX2 enables juvenile-like ocular dominance plasticity in adults. **A,** Adult mice were transfected (p110-118) with AAV2-dnNPTX2. The intrinsic signal responses to stimulation in each eye were recorded in the V1 contralateral to the deprived eye before and after 3 days of monocular deprivation starting at p109-p129. **b,** Example experiment. Left: vasculature pattern of the imaged region used for alignment. Middle: magnitude map of the visual response from the eye contralateral [C] or ipsilateral [I] to the imaged hemisphere. Gray scale (bottom): response magnitude as fractional change in reflection x10^4^. Arrows: L, lateral, R, rostral. Right: histogram of ocular dominance index (ODI) is illustrated in the number of pixels (x-axis: ODI, y-axis: number of pixels). **C-E**, Summary of changes in response amplitude of the deprived contralateral eye (left) and the non-deprived ipsilateral eye (middle), and the change of ODI (right) before (b) and after (a) MD. Thin line: individual experiments; thick line and symbols: average ± s.e.m. For comparison, boxes in **E** depict the 95% CI range of ODI of 15 non-transfected, adult mice (light grey: normal-reared; dark grey: deprived).

## DISCUSSION

Deprivation studies indicate that rapid disinhibition of pyramidal neurons enables plasticity of glutamatergic networks. Here we describe a novel form of synaptic plasticity that mediates cortical disinhibition induced by monocular deprivation: the “all-or-none” elimination of a subset of Pyr➔PV connections. This “all-or-none” elimination of Pyr➔PV inputs is distinct from most other models of synaptic weakening, in that all synaptic contacts between affected pairs of neurons are lost. Pyr➔PV synapse elimination also contrasts with perinatal pruning of exuberant inputs (Chen and Regehr, 2000; Kano and Watanabe, 2019; Patel et al., 2014) in that it is transient, lasting only one day, and it is not accompanied by a compensatory increase in the strength of the remaining inputs. In addition, the elimination of Pyr➔PV inputs is also unique in that it is highly specific, affecting only inputs from nearby layer 2/3 pyramidal neurons, leaving inputs from distant pyramidal neurons in layer 2/3 or in layer 4 unchanged. Finally, the Pyr➔PV input elimination is contingent on a reduced NPTX2 content, and it is a seemingly obligatory step for subsequent changes in ocular dominance.

The observation that the MD-induced change in Pyr➔PV-IN inputs is both all-or-none, and transient, is somewhat counter intuitive, and it is unclear how temporary input elimination best serves ODP. Nevertheless, it seems plausible that the process is initiated by changes in neural activity resulting from diminished visual drive during MD. A reduced correlation of pre-postsynaptic activity in Pyr➔PV pairs after MD is expected to induce associative long-term synaptic depression in these connections (Huang et al., 2013; Lamsa et al., 2010) and may promote synapse elimination as reported in other types of depressed synapses (Henson et al., 2017; Oh et al., 2013). We propose that the reduction in the availability of NPTX2 induced by MD contributes to the destabilization of depressed Pyr➔PV inputs in V1. Notably, associative LTD in Pyr➔PV connections and synapse elimination in CA1 pyramidal cells require mGluR5 activation (Huang et al., 2013; Wilkerson et al., 2014), raising the possibility that the elimination of individual synapses and cell-to-cell connections share common initial mechanisms. The elimination of Pyr➔PV inputs is highly local, occurring only in layer 2/3 Pyr➔PV pairs separated by less than ~25 μm. This spatial requirement matches the diameter of recently reported microcolumns in mouse cortex, suggesting that the distribution of disconnected synapses might reflect the co-regulation of locally connected neuronal networks (Maruoka et al., 2017). The preferential local elimination might also be attributed to cortical retinotopy, as the decrease in correlated activity during MD might be more pronounced in proximal than distal pairs. The absence of an overt change in connectivity between layer 4 Pyrs➔layer 2/3 PVs following MD may be due to low expression of NPTX2 in layer 4 neurons (Allen brain Mouse Atlas). Finally, the loss of Pyr➔PV inputs might be transient because subsequent homeostatic changes in glutamatergic synapses might restore patterns of neural activity conducive to reestablish connectivity (Bridi et al., 2018; Hengen et al., 2013; Keck et al., 2017; Lee and Kirkwood, 2019).

Previous studies, including ours (Gu et al., 2016; Sun et al., 2016) have implicated the growth factor NRG1 in plastic changes induced my MD. In contrast, here we show that application of exogenous NRG1 neither prevented nor restored the transient disconnection of MD-induced local Pyr➔PV inputs in layer 2/3, arguing against a direct role of NRG1 in this process. A recent report utilizing a glutamate uncaging approach reported that applied NRG1 can restore a widespread reduction of excitatory input onto layer 2/3 PVs induced by MD(1) (Sun et al., 2016). However, the uncaging approach can recruit and record responses with a large polysynaptic component and the paired whole cell recordings used here may be preferable for resolving local monosynaptic connections. An attractive and simple possibility to reconcile these apparent discrepancies is that MD induces a NPTX-dependent disconnection of local Pyr-PV inputs in layer 2/3, and a NRG1-dependent reduction in local excitatory inputs onto layer 4 PVs. This would be in line with the notion that distinct mechanisms govern synaptic plasticity in different cortical layers (Fong et al., 2020; Frantz et al., 2020; Jiang et al., 2007), and is consistent with our observation that NRG1 affects excitatory inputs onto layer 2/3 PVs originating from layer 4, but not layer 2/3. If correct this scenario allows for two molecularly distinct mechanisms to regulate excitatory inputs onto PVs, that are anatomically segregated, thereby minimizing functional redundancy.

The rapid and transient elimination of approximately 50% of the local excitatory inputs onto PVs is likely a major determinant of the deprivation-induced cortical disinhibition attributed to the reduced activity of PVs (Kuhlman et al., 2013; Ma et al., 2019). However, other synaptic mechanisms might also contribute. For example, even if the strength and connectivity of the PV output is not affected by MD(1) (supplementary Fig. 2), changes in short-term synaptic dynamics, normally tuned for high frequency transmission (Bridi et al., 2020), could enhance cortical activity. Similarly, changes in the electrical coupling between PVs, and tonic inhibition, known to be affected by deprivation in pyramidal neurons (Huang et al., 2015), could enhance cortical activity. On the other hand, unlike the case of barrel cortex (Gainey et al., 2018) changes in the intrinsic excitability of layer2/3 PVs is not a factor for disinhibition (Sun et al., 2016). On the other hand, unlike the case of barrel cortex (Gainey et al., 2018) changes in the intrinsic excitability of layer2/3 PVs is not factor for disinhibition (Sun et al., 2016).

The critical periods for ocular dominance plasticity (Huang et al., 2015; Huang et al., 2010) and the plasticity of Pyr➔PV-IN connectivity are highly coincidental, both terminating by p90. Furthermore, manipulations of NPTX2 function that prevent and promote the transient disconnection of Pyr➔PV-IN inputs also prevent/promote ocular dominance shift after MD. We therefore surmise that the transient elimination of local Pyr➔PV inputs is an obligatory initial step for the subsequent remodeling of glutamatergic circuitry underlying ocular dominance plasticity.

An attractive candidate step to follow the loss of Pyr➔PV connections could a rapid increase in the proportion of synaptic GluN2b over GluN2a subunit, which is observed after brief visual deprivation (Bridi et al., 2018; Guo et al., 2012). The change in NMDAR composition lowers the threshold for the induction of synaptic plasticity, including LTP and LTD elicited with spike-timing protocols, and is consequence of cortical disinhibition (Bridi et al., 2018; Guo et al., 2012). In this scenario, glutamatergic connectivity would not be intrinsically more plastic during the critical period, but plasticity is enabled by the change in GluN2b/GluN2a following the local elimination of Pyr➔PV inputs. The termination of the critical period for ODP, would therefore be a direct consequence of the loss of plasticity of the Pyr➔PV inputs. In this general context, it will be of interest to determine whether NPTX2-dependent disconnection of Pyr➔PV inputs also regulates other candidate mechanisms of ODP, like the abundance of silent synapses (Xu et al., 2020). Finally, it will be also of interest to examine whether disconnection of local Pyr➔PV inputs contributes to the changes in cortical criticality after MD (Ma et al., 2019) and to pathology in neurological conditions known to exhibit marked alterations in NPTX2 abundance, including Alzheimer disease and schizophrenia (Xiao et al., 2017).

## Author Contributions

D.S, S.Y.H and M.C.D.B. collected and analyzed slice electrophysiology data. S.H., S-Y.R., I.H. and RH collected, analyzed and interpreted the imaging data, J.Z made the NPTX2 viruses, S.M, S.R., R.H., M.G and E.M.Q collected, analyzed and interpreted the immunohistology and immunochemistry data. D.S., P.W., E.M.Q. and A.K. wrote the manuscript.

## Acknowledgements

We thank Prof. HK Lee for valuable advice on the design and writing and Prof. S. Zeger for statistical advice. Supported by grants R35 NS097966 to PW, PO1 AG009973 to M.G., R01EY016431 to EMQ, R01EY12124 to A.K., R01EY025922 to EMQ and AK

## STAR Methods

### RESOURCE AVAILABILITY

#### Lead Contact

Further information and requests for resources and reagents should be directed to and will be fulfilled by the lead contact, Alfredo Kirkwood.

#### Materials Availability

This study did not generate new unique reagents.

#### Data and Code Availability

Physiological or imaging source data are available from Mendeley Data or from the corresponding author on request. No further unique datasets or codes were generated in this study.

### EXPERIMENTAL MODEL AND SUBJECT DETAILS

#### Animals

C57BL/6, PV-cre;Ai14, Scnn1a-Tg3-cre;Ai32;PV-tdT, and Scnn1a-Tg3-cre;G42 mice of both sexes were used. All subjects were group-housed (5 or fewer per cage) on a 12 hr light/dark cycle, with food and water available ad libitum. All protocols and procedures conformed to the guidelines of the US Department of Health and Human Services Office of Laboratory Animal Welfare (OLAW) and were approved by the Institutional Animal Care and Use Committees of the University of Maryland, Johns Hopkins University, or both. These committees followed the guidelines established by the Animal Care Act and National Institutes of Health (NIH). For specific optogenetic stimulation in layer 4, we developed a triple transgenic line crossing the L4-Cre mouse line with the Ai32 mouse line (Madisen et al., 2012) that has a conditional allele of Rosa-CAG-LSL-ChR2(H134R)-EYFP-WPRE to drive ChR2/EYFP fusion protein expression in L4 principal cells. We then crossed L4-cre positive and Ai32 homozygous mice with PV-tdT mice.

### METHOD DETAILS

#### Monocular deprivation

Naive mice were monocularly deprived of vision for 1, 2, or 3 d beginning at postnatal day 21–25, 27–31, 50–54, or 100–118. Mice were age-matched across groups. Monocular deprivation (lid suture) was performed under isoflurane anesthesia (2-3% for induction; 1.5% for maintenance). The margins of the upper and lower lids of one eye were trimmed and sutured together. Neosporin was applied to the sutured eye. Animals were disqualified in the event of suture opening or infection.

#### Drug administration

The recombinant human NRG1-β1/HRG1-β1EGF domain (7.5 kDa; R&D Systems, Minneapolis, MN), corresponding to residues 176–246 of the soluble EGF domain of neuregulin β1 common to all NRG1 splice variants, was dissolved in sterile saline. We used residues 177-241 from the same splice variant in some cases, with no difference in results between the two peptides (7.5 kDa; PROSPEC Protein Specialists, East Brunswick, NJ). Monocularly deprived mice were subcutaneously administered with two injections of recombinant NRG1 (1 μg NRG1 in 50μL of sterile saline per injection per mouse). The first injection was applied immediately before lid suture, and the second, one hour before the animal was sacrificed for electrophysiological experiments.

#### Viral injections

Mice were anesthetized and head-fixed in a stereotaxic device (Kopf Instruments, Los Angeles, CA) under 1.5%–2% isoflurane. V1 was located in the left hemisphere using stereotaxic coordinates (3.6 mm posterior, 2.5 mm lateral to bregma). When the bregma-lambda distance was different than 4.2 mm, the stereotaxic coordinates were scaled. A craniotomy (~0.5 mm) was made, and the virus was injected (600 nL; 100 nL/sec) into layer 2/3 or layer 4 (0.3 or 0.4 mm from the cortical surface, respectively) 10-35 d before experimentation. For 2P imaging, during craniotomy AAV2.CaMKII.NPTX2-SEP was injected into three sites (100 nL per site; 5 nL/sec). Viral injection of AAV2.CamKII.NPTX2-SEP or AAV2.CaMKII.GFP was done by bulk regional viral injection to the visual cortical area of neonatal PV-Cre;Ai14 mice at p0–2. Briefly, neonatal mice were cryo-anesthetized (Phifer and Terry, 1986) in an ice-cold chamber and positioned dorsal side up and secured with an adhesive bandage across the upper body. The visual cortical area was targeted with anatomical landmarks, including occipital fontanelle and lambdoid suture, visible through the neonatal skin. 600 nL of AAVs were injected right beneath the skull with a syringe infusion pump (50 nL/sec). After the injection, mice were kept on a heating pad and returned to their home cage once recovered. For optical imaging of intrinsic signaling, AAVs were injected at three locations targeting layer 2/3 of the binocular region of the V1 (500 nL/each injection site). For 2P imaging, during craniotomy AAV2-CaMKII-NPTX2-SEP was injected into three sites (100 nL per site) using a glass pipette (pulled by pipette puller, Narishige, Japan) and Nanoject (Drummond Scientific Company, PA) at a rate of 5 nL / second.

### SLICE ELECTROPHYSIOLOGY

Visual cortical slices were prepared as previously described (Huang et al., 2012). Each mouse was anesthetized using isoflurane vapors. Mice older than p50 were transcardially perfused with ice-cold dissection buffer containing (in mM): 212.7 sucrose, 5 KCl, 1.25 NaH2PO4, 10 MgCl2, 0.5 CaCl2, 26 NaHCO3, and 10 dextrose, bubbled with 95% O2 / 5% CO2 (pH 7.4)). Then mice were immediately decapitated, the brain was removed, and slices 300 μm thick were cut in icecold dissection buffer. The slices were transferred to artificial cerebrospinal fluid (ACSF), incubated at 30°C for 30 minutes, and then kept at room temperature for 30 minutes until they were transferred to the recording chamber. ACSF was similar to dissection buffer except that sucrose was replaced by 124 mM NaCl, MgCl2 was lowered to 1 mM, and CaCl2 was raised to 2 mM. Visualized whole-cell recordings were made from FS(PV)-INs and pyramidal neurons with glass pipettes filled with 130 mM K-gluconate, 10 mM KCl, 10 mM HEPES, 0.2 mM EGTA, 0.5 mM Na3GTP, 4 mM MgATP, and 10 mM Na-Phosphocreatine (pH 7.2–7.3, 280–290 mOsm). Only cells with series resistance <25 MΩ (with <20% variation over the experiment) were included. Data were filtered at 4 kHz and digitized at 10 kHz using Igor Pro (Wave Metrics). The intersomatic distance was calculated as the Euclidean distance between the centers of both somata. Unitary excitatory postsynaptic potentials (uEPSCs) were recorded in voltage-clamp in the PV-INs at −70 mV and evoked by suprathreshold somatic current injection (2 ms) in presynaptic pyramidal neurons. uIPSCs were recorded in voltage-clamp in pyramidal neurons at 0 mV and evoked by suprathreshold somatic current injection (2 ms) in presynaptic PVs (Jiang et al., 2010). For presynaptic parameters, all presynaptic cells were depolarized with 1.7pA current injection. At least 20 responses evoked at 0.1 Hz with paired-pulse stimulation (interstimulus interval: 50 ms for Pyr→PV pairs; 100 ms for PV→Pyr pairs) were used to confirm a synaptic connection and to compute the amplitudes of the unitary responses. When the temporal course of evoked EPSCs was measured (Fig. 4b,c), recordings were excluded if baselines were not stable.

#### Mean-variance and presynaptic analysis

were performed on responses evoked by at least 15 stimuli trains delivered at 50 Hz and 20 s intervals. The uEPSC amplitude was measured for each stimulus, and the mean and variance were plotted against each other. Quantal size (q) was obtained from the linear regression slope for all events, excluding the two events with the highest average release probability (Clements and Silver, 2000). The Schneggenburger-Meyer-Neher (SMN) approach (Schneggenburger et al., 1999) was used to explore whether the readily releasable pool (RRP) of presynaptic vesicles was affected. The cumulative amplitude was plotted versus the stimulus number, and a linear regression was adjusted to the last five stimuli. In this model, the RRP replenishment rate is represented by the linear fit slope and the RRP size by the y axis intercept. We considered only those cases in which the R^2^ value of the fit was >0.95 for presynaptic analysis and >0.5 for mean-variance analysis.

#### Maximal postsynaptic currents (PSC max)

recorded in PV-IN were evoked every 15 seconds by electrical stimulation using a theta glass bipolar electrode placed in the middle of the cortical thickness. We used a Cs-gluconate based internal solution containing (in mM) 120 Cs-gluconate, 8 KCl, 10 HEPES, 1.12 EGTA, 0.5 Na3GTP, 4 Na2ATP, 10 Na-Phosphocreatine, and 5 Lidocaine; pH 7.2–7.3, 280–290 mOsm). Excitatory and Inhibitory PSC max (EPSC and IPSC max) were recorded in Voltage clamp at −40 and −70 mV, respectively. For IPSC max measurements, 25 μM 6-cyano-7-nitroquinoxaline-2,3-dione (CNQX) and 100 μM DL-2-amino-5 phosphonopentanoic acid (DL-APV) were included in the bath, and electrode capacitance and series resistance were compensated (Prediction % 90, Correction% 80-85, bandwidth 3 kHz).

#### NMDA/AMPA ratios

were calculated from excitatory currents onto PV-IN evoked from layer 2/3 by electrical stimulation. 2 times the stimulus intensity that evoked the minimal AMPA response was used. An internal solution containing (in mM) 102 cesium gluconate, 5 TEA chloride, 3.7 NaCl, 20 HEPES, 0.3 Na-GTP, 4 Mg-ATP, 0.2 EGTA, 10 BAPTA, 5 QX-314 (pH 7.2, ~300 mOsm) under voltage clamp (Vh = −70 mV for AMPA and Vh = + 40 mV for NMDA). To isolate glutamate evoked currents and minimize multisynaptic responses, ACSF in the recording chamber contained 2.5 μM gabazine, 25 μM CNQX, 10 μM adenosine, 4 mM CaCl2, and 4 mM MgCl2.

#### PV/Pyr EPSC response ratios

were calculated from L4-evoked excitation onto L2/3 PV-IN and neighboring pyramidal cell (12-24 μm apart). An input-output curve for EPSPs onto each cell was measured by simultaneous pair recordings using identical light intensity. The stimulation threshold was defined as the light intensity that elicited the minimal PV response with no failures.

### IMMUNOHISTOFLUORESCENCE

Subjects were anesthetized with isoflurane and perfused with phosphate-buffered saline (PBS) followed by 4% paraformaldehyde (PFA) in PBS. Brain was post-fixed in 4% PFA for 24 hours followed by 30% sucrose for 24 hours, and cryoprotectant solution (0.58 M sucrose, 30% (v/v) ethylene glycol, 3 mM sodium azide, 0.64 M sodium phosphate, pH 7.4) for 24 hr. Coronal sections (40 μm) were made on a Leica freezing microtome (Model SM 2000R). Sections were blocked with 4% normal goat serum (NGS) in 1X PBS for 1 hour. Antibodies were presented in a blocking solution for 18 hours, followed by appropriate secondary antibodies.

#### Antibodies

The following antibodies/dilutions were used: mouse anti-parvalbumin (PV, Millipore) RRID:AB_2174013, 1:2000; guinea pig anti-VGluT1 (Millipore) RRID:AB_2301751, 1:2000; followed by appropriate secondary IgG conjugated to Alexa-488, 546 (**Thermo Fisher Scientific**) RRID:AB_2534089, RRID:AB_2534093, RRID:AB_2535805, 1:1000.

#### Confocal imaging and analysis

Images were acquired on a Zeiss LSM 710 confocal microscope. A maximal intensity projection of a z-stack (11 slices x 0.5 μm images) was acquired at 40X (Zeiss Plan-neofluar 40x/1.3 Oil DIC, NA=1.3). PV+ somata were identified by size exclusion (20-200 mm^2^) and fluorescence intensity (auto threshold + 25). Co-localization of VGluT1 puncta on PV somata were analyzed in single Z-section images taken at 40X, using Fiji. After the threshold function was applied to VGluT1 puncta (autothreshold+25), co-localized puncta were identified by size exclusion (0.1 μm2 < 2.0 μm2).

### IMMUNOCIYTOCHEMISTRY

#### Western blots

Protein from the dissected visual cortex was extracted with RIPA buffer such that the non-deprived hemisphere was used as a control for each deprived side of the mouse. 12ug of protein were run on 4-12% Nupage gels and transferred to nitrocellulose membrane. Membranes were probed Nptx2 (1:1000; Abcam: ab191563) and visualized a Chicken anti-Rabbit IgG, Alexa Fluor 647 secondary antibody (Thermo Fisher Scientific: A-21443). Tubulin was used as a loading control (α-Tubulin Mouse mAb; Cell Signaling: cat# 3873). Fluorescent bands were visualized on Typhoon 9410 variable mode imager (Amersham) with settings optimized for linear detection of Nptx2 and tubulin band intensities. Scans were quantified using ImageQuant with a fixed area for each protein. Background subtracted average intensities were z-scored to combine across gels.

### IMAGING

#### Craniotomy

To install the glass window, juvenile mice (p27-31) were first anesthetized with isoflurane (2.5% for induction; 1.5% for maintenance in oxygen). After removal of the skin, connective tissue was gently removed by the blade with hydroperoxide. Translucent dental cement (C&B metabond, Parkell, NY) was applied to the exposed dry skull region. A 3 mm diameter craniotomy was made with a No.11 surgical blade over the putative visual cortical area, and the virus injection was made when corresponded. Three-layered (one 5 mm, two 3 mm glass cover glass) glass window was installed to the craniotomy, and dental cement was applied to fix the window. For 2-photon microscopy, the custom-made metal head bar was attached using the dental cement Superbond (Sun Medical, Japan). For the older animals (~p90), a craniotomy was performed with a dental drill. Atropine (0.05 mg/kg, s.c.) and dexamethasone (4.8 mg/kg, i.m.) were injected to reduce mucosal secretion and brain edema, respectively.

#### Optical imaging of the intrinsic signal

was performed with the mice anesthetized with isoflurane (2-3% for induction; 0.7-1.2% for maintenance in oxygen) supplemented with chlorprothixene (2 mg/kg, i.p.). The mice were placed with head-fixed in front of the LCD monitor for visual stimulus. Atropine was injected subcutaneously to reduce mucosal secretion (0.05 mg/kg). Eye drops were administered to keep eyes moist, and body temperature was maintained at 37°C with a heating pad and a rectal probe. Heart rate was monitored throughout the experiment by electrocardiogram. Acquisition of intrinsic signal was performed following the method optimized to measure the ocular dominance in mice (Cang et al., 2005; Kalatsky and Stryker, 2003). Briefly, visual stimulation evoked intrinsic signals were acquired using a Dalsa 1M30 CCD camera (Dalsa, Waterloo, Canada) controlled by custom software. The surface vasculature and EYFP fluorescence were visualized with 555 nm LED illumination, and the intrinsic signals were imaged with 610 nm LED illumination. For the acquisition of the intrinsic signal, the camera was focused 600 mm below the surface of the skull. An additional red filter was interposed to the CCD camera, and intrinsic signal images were acquired. A high refresh rate monitor (1024×768 @ 120 Hz; ViewSonic, Brea, CA) was placed 25-cm in front of the mouse, with their midline aligned to the midline of the mouse for visual stimulus. The visual stimulus presented was restricted to the binocular visual field (−5° to +15° azimuth) and consisted of a thin horizontal bar (height = 2°, width = 20°) continuously presented for 5 minutes in upward (90°) and downward (270°) directions to each eye separately. The cortical response at the stimulus frequency was extracted by Fourier analysis. The two maps generated with the opposite direction of the drifting bar were averaged for each eye to calculate the response amplitude and the ocular dominance index (ODI). The ODI was computed as follows: (1) the intensity maps were smoothed by 5×5 low-pass Gaussian filter; (2) the binocular region of interest (ROI) was defined at 30% of peak response amplitude of the smoothed intensity map from the ipsilateral eye; (3) the response amplitude of each eye was calculated by averaging the intensity of all pixels in the ROI; (4) the ODI was calculated by the average of (C-I)/(C+I) of all pixels in the ROI where C and I are the contralateral (C) and ipsilateral (I) eye respectively.

#### In vivo 2-photon awake microscopy for NPTX2-SEP imaging

Confocal microscopy was performed with a laser scanning microscope (Olympus, Japan) equipped with an ultra-sensitive GaAsP detector (Hamamatsu, Japan) and a Galvanometer scanner (Thorlabs, NJ). 2-photon excitation was carried out by ultrafast Ti:Sapphire laser Mai Tai eHP DeepSee (Spectra-Physics, CA), operating the wavelength at 920 nm for visualizing NPTX2-SEP with GFP filter using 20X objective XLUMPLFL20XW (Olympus, Japan). Hardware operation and image acquisition were performed with PrairieView software (Bruker, MA). 11-13 days after surgery, mice were handled for 10 to 15 min and acclimated to the imaging setup for 15 to 30 mins by fixing their head bars to the custom-made head fixation apparatus and imaging platform of a cylindrical treadmill. Intrinsic signal imaging (ISI) was performed to identify the primary visual cortex (V1). On day 14, the treadmill was fixed to minimize the motion-related artifacts, and 3D z-stack images were acquired at V1 and non-V1 regions. 512 by 512 resolution images with 8 averages were acquired from 0 to 180 μm from dura in an interval of 5 μm (total 37 image stacks), where lower 100 μm was selected for layer 2/3 analysis. The images were acquired in the same location with the same imaging parameters for longitudinal study with MD in a reference of blood vessel morphology captured by epifluorescence imaging with 605 nm LED excitation (Thorlabs, NJ). To study the effect of MD on NPTX2 secretion, the control image was acquired before the contralateral monocular eye suture, followed by imaging for 3 consecutive days in the same location.

#### NPTX2-SEP spot 3D analysis

All 3D z-stack time-series (4D) images were preprocessed with ImageJ (FIJI) and analyzed with Imaris software (Bitplane, Zurich, Switzerland). Each set of 3D z-stack images was concatenated to generate 4D images, background-corrected by ‘bleach correction’ using histogram matching algorithm, and 3D-drift corrected by ‘Correct 3D drift’ algorithm (Parslow et al., 2014) (FIJI). The xyz-corrected portion of the processed 4D images was selected for NPTX2-SEP spot analysis with Imaris software. The NPTX2-SEP spots were detected with the ‘surface detection’ function, maximum diameters of spheres detected to be 3 μm, and corresponding values within the 3D area were extracted (spot number, intensity, and volume). The ‘total volume’ is defined by the sum of 3D compartments occupied by NPTX2-SEP positive voxels. The spot number and the total volume were normalized with the median value of the ‘before’ condition and presented per each animal. The mean intensity and mean volume were normalized by the median value of ‘before’ condition, and all the spots from all animals were calculated. (The histogram analysis was done with 5 and 25 bin for intensity and volume, respectively (Prism Graphpad).)

### QUANTIFICATION AND STATISTICAL ANALYSIS

Sample sizes were chosen to correspond with previous studies in which the effects of visual manipulation were measured. Animals were randomly placed into experimental groups. Whenever was possible, treatments were applied in the same animal (e.g. deprived hemisphere versus non-deprived hemisphere) or in the same cell (e.g. electrical stimulation from layer 4 or layer 2/3, Fig 4b). For puncta quantification (Fig. 3c,d) and western blots (Supplementary Fig. 4), the experimenter was blind to the treatment of samples during analyses the analysis. Exclusion criteria are described for each experiment. Normality was determined using the D’Agostino test, and variance was compared using Levene’s median test. Groups with normally distributed data were compared using two-tailed paired or unpaired t-tests, one-way ANOVAs, or one-way repeated-measures ANOVAs, as indicated. Holm–Sidak post hoc tests were used for multiple comparisons following one-way ANOVAs. Groups that were not normally distributed were compared using nonparametric Wilcoxon matched-pairs signed-rank test, Mann-Whitney test, or ANOVAs on ranks (followed by Dunn’s post hoc test for multiple comparisons). Statistical outliers were detected using pre-established criteria (ROUT test) and excluded from the analysis. Data are presented as averages ± s.e.m. Statistical analyses were performed using GraphPad Software (Prism, San Diego, CA).

### KEY RESOURCES TABLE

| REAGENT or RESOURCE | SOURCE | IDENTIFIER |
| --- | --- | --- |
| **Antibodies** |  |  |
| mouse anti-parvalbumin | Millipore | RRID:AB_2174013 |
| guinea pig anti-VGluT1 | Millipore | RRID:AB_2301751 |
| Goat anti-Mouse, Alexa Fluor 546 | Thermo Fisher Scientific | Cat# A-11030, RRID:AB_2534089 |
| Goat anti-Rabbit, Alexa Fluor 546 | Thermo Fisher Scientific | Cat# A-11035, RRID:AB_2534093 |
| Goat anti-Mouse, Alexa Fluor 647 | Thermo Fisher Scientific | Cat# A-21236 RRID:AB_2535805 |
| α-Tubulin Mouse mAb | Cell Signaling | Cat # 3873 |
| Rabbit anti-mouse NPTX2 | abcam | abcam: ab191563 |
| Chicken anti-Rabbit IgG, Alexa Fluor 647 | Thermo Fisher Scientific | Cat# A-21443, RRID:AB_2535861 |
| **Bacterial and Virus Strains** |  |  |
| AAV2/2.CaMKII.dnNPTX2-EGFP | O'Brien et. Al. 2002 | N/A |
| AAV2.CaMKII.NPTX2-SEP | PENN vector core | https://pennvectorcore.med.upenn.edu/ |
| AAV2.CaMKII.GFP | UNC vector core | https://www.med.unc.edu/genetherapy/vectorcore/in-stock-aav-vectors/boyden/ |
| AAV9.Syn.Flex.ChrimsonR-tdT | Klapoetke et al. 2014 | RRID:Addgene_62723 |
| Chemicals, Peptides, and Recombinant Proteins |  |  |
| tetrodotoxin citrate | Abcam | Cat # AB120055 |
| DL-AP5 | Tocris | Cat # 0105 |
| Adenosine 5'-triphosphate magnesium salt | Sigma Aldrich | Cat # A9187 |
| Guanosine 5' Triphosphate Sodium Salt Hydrate | Sigma Aldrich | Cat # G8877 |
| Phosphocreatine disodium salt hydrate | Sigma Aldrich | Cat # P7936 |
| CNQX disodium salt hydrate | Sigma Aldrich | Cat # C239 |
| SR-95531 (gabazine) | Sigma Aldrich | Cat # S106 |
| Human NRG1-β | R&D systems | Cat # 396-HB-050 |
| Human NRG1-β | PROSPEC | Cat # CYT-733 |
| Lidocaine N-ethyl Bromide (QX-314) | Sigma Aldrich | Cat # L5783 |
| Phosphocreatine disodium salt hydrate | Sigma Aldrich | Cat # P7936 |
| Guanosine 5' Triphosphate Sodium Salt Hydrate | Sigma Aldrich | Cat # G8877 |
| Adenosine 5'-triphosphate magnesium salt | Sigma Aldrich | Cat # A9187 |
| Adenosine | Sigma Aldrich | Cat # 01890 |
| Cesium Hydroxide | MP Biomedicals | Cat # 101328 |
| 4-(2-Hydroxyethyl)-1-Piperazineethanesulfonic Acid (HEPES) | Fisher bioreagents | Cat # BP310-100 |
| Sodium Chloride | Fisher Chemical | Cat # S271 |
| Dextrose Anhydrous | Fisher Chemical | Cat # D16 |
| 1,2-Bis(2-Aminophenoxy)ethane-N,N,N',N'-tetraacetic acid (BAPTA) | Sigma Aldrich | Cat # A4926 |
| Ethylene glycol-bis(2-amino-ethylether)-N,N,N',N'-tetraacetic acid (EGTA) | Sigma Aldrich | Cat # E4378 |
| Potassium D-gluconate | Sigma Aldrich | Cat # G4500 |
| Potassium chloride | Fisher Chemical | Cat # P217 |
| Tetraethylammonium chloride | Sigma Aldrich | Cat # T2265 |
| Calcium Chloride Dihydrate | Fisher Chemical | Cat # C79 |
| Magnesium Chloride Hexahydrate | Fisher Chemical | Cat # M33 |
| D-Gluconic acid | Sigma Aldrich | Cat # G1951 |
| D-Sucrose | Fisher bioreagents | Cat # BP220 |
| Deposited Data |  |  |
| Raw and analyzed data | This paper and Mendeley DATA |  |
| Experimental Models: Organisms/Strains |  |  |
| B6 mouse: C57BL/6J | The Jackson Laboratory | RRID:IMSR_JAX:000664 |
| L4-cre mouse: B6;C3- Tg(Scnn1a-cre)3Aibs/J | The Jackson Laboratory | RRID:IMSR_JAX:009613 |
| Ai32 mouse: B6;129S-Gt(ROSA)26Sor<sup>tm32(CAG-COP4<sub>H134R/EYFP</sub>)Hze/J</sup> | The Jackson Laboratory | RRID:IMSR_JAX:012569 |
| Ai14 mouse: B6.Cg-Gt(ROSA)26Sor<sup>tm14(CAG-tdTomato)Hze/J</sup> | The Jackson Laboratory | RRID:IMSR_JAX:007914 |
| PV-cre mouse: B6;129P2-Pvalbtm1(cre)Arbr/J | The Jackson Laboratory | RRID:IMSR_JAX:008069 |
| PV-tdTomato mouse: C57BL/6-Tg(Pvalb-tdTomato)15Gfng/J | The Jackson Laboratory | RRID:IMSR_JAX:027395 |
| PV-GFP mouse: CB6-Tg(Gad1-EGFP)G42Zjh/J | The Jackson Laboratory | RRID:IMSR_JAX:007677 |
| Software and Algorithms |  |  |
| Prism | GraphPad | RRID:SCR_002798 |
| IgorPro | Wavemetrics | RRID:SCR_000325 |
| Imaris | Oxford Instruments | RRID:SCR_007370 |
| ImageJ | NIH | RRID:SCR_003070 |
| ImageQuant TL | Cytiva | RRID:SCR_018374 |
| Black Zen software | Zeiss | RRID:SCR_018163 |

