## supplementary figures for "All-or-none disconnection of pyramidal inputs onto parvalbumin-positive interneurons gates ocular dominance plasticity"

Supplementary figure 1 (Figure 1 related)

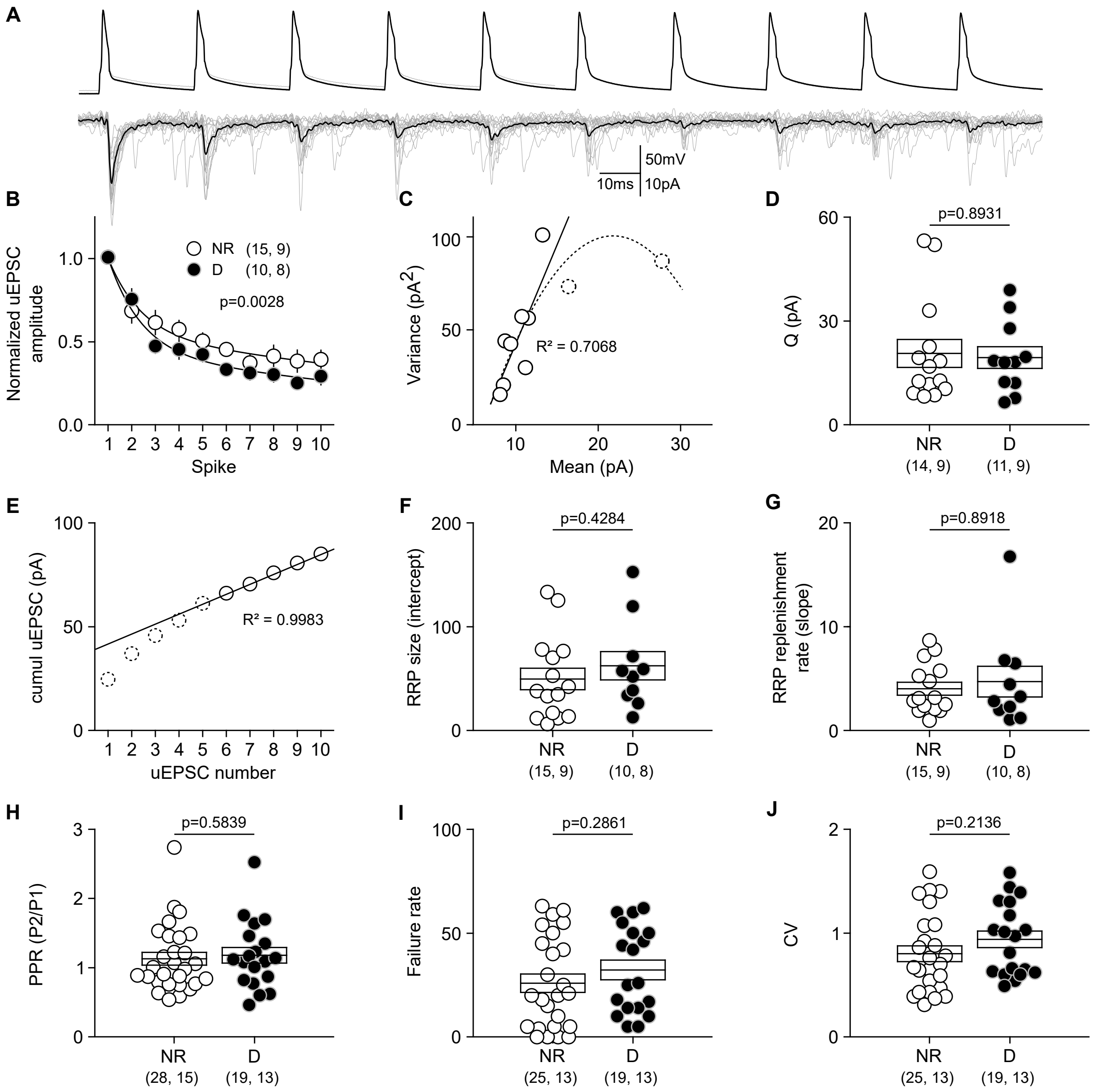

Supplementary Figure 2 (Figure 2 related)

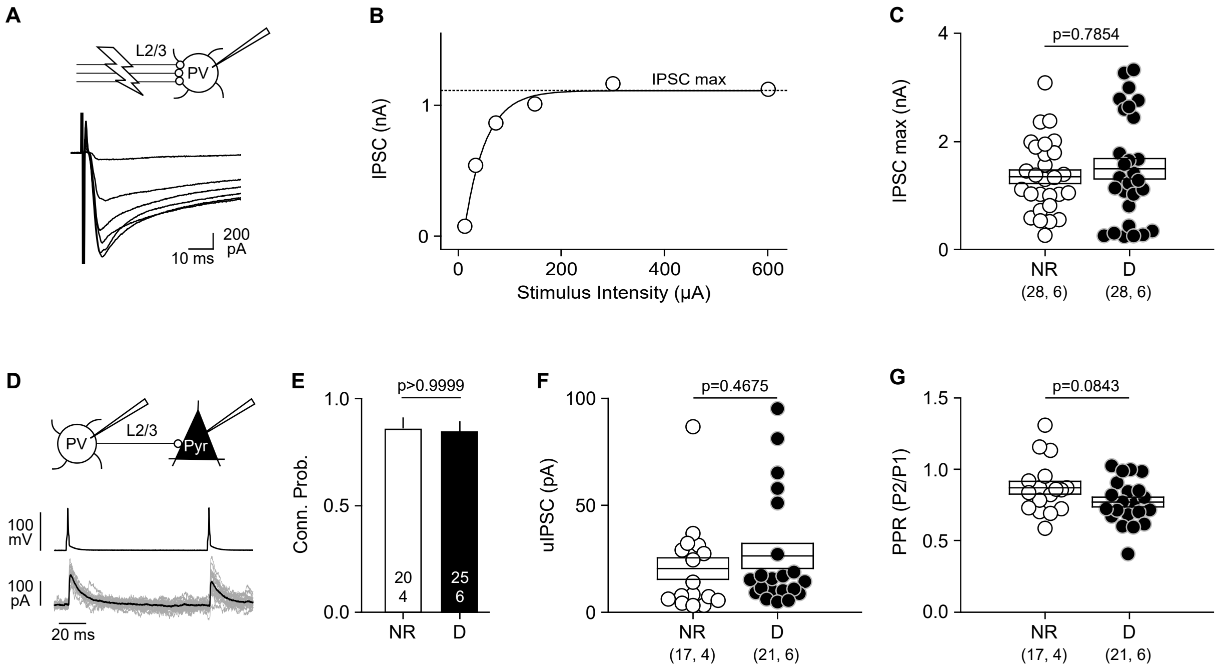

Supplementary Figure 3 (Figure 5 related)

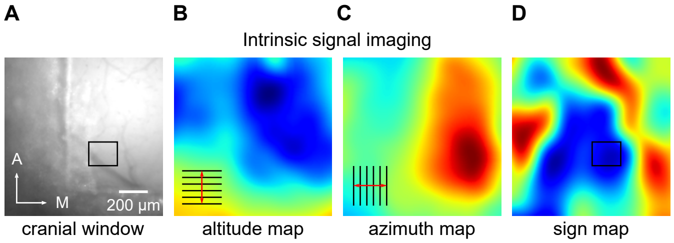

Supplementary Figure 4 (Figure 5 related)

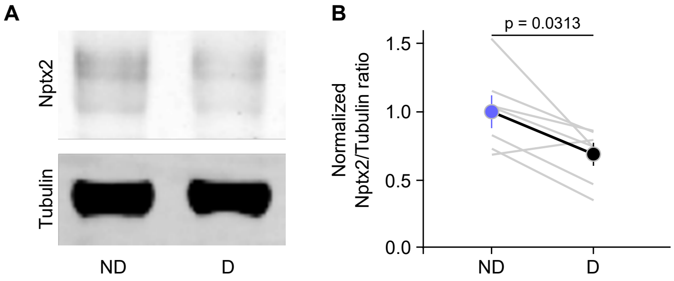

Supplementary Figure 5 (Figure 6 related)

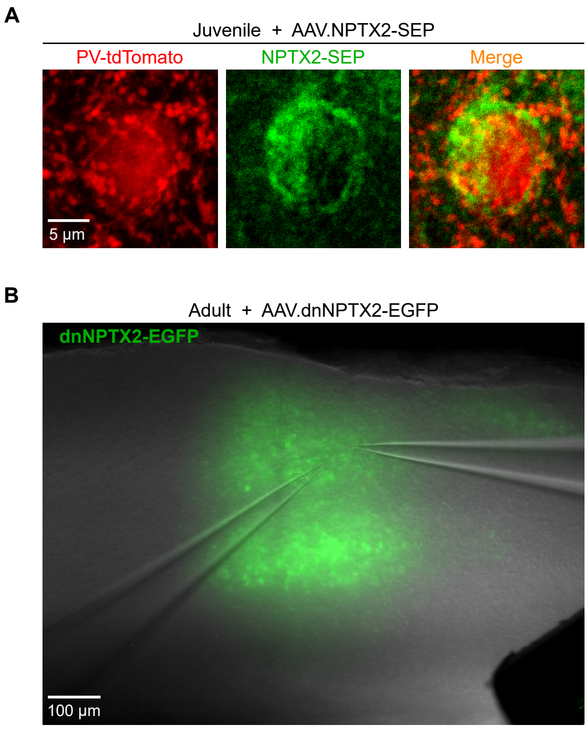

**Supplementary Fig. 1 | MD(1) does not affect parameters of remaining Pyr->PV synapses.** Synaptic parameters were estimated from the responses to 50 Hz trains (10 pulses). **A,** Superimposed traces show an example of 15 consecutive trials (gray) along with the averaged response (black). Scale bars: 50 mV, 10 pA, 20 ms. **B**, The attenuation of the responses (normalized to the first response) during the train was comparable in pairs from NR (white circles) and D (black circles) mice (two-way: ANOVA F[1, 230]=9.097, p=0.0028; interaction F[9, 230]=0.65, p=0.757). **c**, The example illustrates how the quantal size (Q), determined from the initial slope of a plot of mean uEPSC amplitude versus variance as in (ref)(see methods for more details). **D**, Q was not different in pairs from NR and D mice (MW-test U=74, p=0.893) **E**, the example illustrates how the linear relationship between the cumulative uEPSC amplitude vs stimulus number was used to estimate the readily releasable pool (RRP) size (intercept) and replenishment rate (slope) (Schneggenburger et al., 1999) see methods for details. **F**,**G,** MD(1) did not affect either the RRP size (**F** : MW-test U=60, p=0.4284) or the RRP replenishment rate (**G**, MW-test U=72, p=0.892). In addition, MD(1) did not affect the paired-pulse ratio (P2/P1)of the responses (**H:** MW-test U=240, p=0.583), the uEPSC failure rate (**i**: MW-test U=192, p=0.2861) or the coefficient of variation (CV) of the responses (**j**: MW-test U=185, p=0.2136). The number of cells or cell pairs and mice is indicated in parenthesis in **B-F.**

**Supplementary Fig. 2 | does not affect the synaptic output of PV-Ins. A-C,** Maximal IPSCs (IPSCmax) in L2/3 PV-INs. **A**, IPSCs were evoked by electrical stimulation delivered ~100um laterally. **B**, the IPSCmax were determined from the responses to stimuli series of increasing intensity. **C**, IPSCmax were not different in PV-INs from NR and D mice (MW-test U=375, p=0.7854). **D**-**G**, MD(1) does not affect the PV-IN→Pyr connectivity. **D**, shows an example of uIPSCs (individuals: gray; average: black) evoked in a pyramidal cell (Pyr) by PV-IN firing. **E**, comparable connection probability in pairs from NR and D mice (F-test p>0.9999). **F**, uIPSC amplitude in connected pairs from NR and D mice (MW-test U=153, p=0.4675). **G**, uIPSC paired pulse ratio (PPR: 100 ms interval) in connected pairs from NR and D mice (t-test F[31.5]=1.782, p=0.0843).

**Supplementary Fig. 3 | Localization of V1 with intrinsic signal imaging for the analysis of NPTX2-SEP. A**, The epifluorescence image of the cranial window. The black box area is where two-photon microscopy was performed. **B-D**, Altitude, azimuth, and sign map of the visual responses reveal the V1 region. The grids and arrows in B,C indicate the direction of visual stimulation.

**Supplementary Fig. 4 | Reduced NPTX2 protein content in V1 after MD(1).** Western blot analysis of NPTX2 expression in the non-deprived (ND: open circle) and deprived (D: black circles) V1 of mice after 1 day of MD. NPTX2 levels were relativized to β-actin through densitometry analysis. Left, example experiment. Right; the grey lines connect data of individual mice Individual. Circles connected by the thick line represent mean ± SEM. Wilcoxon test W=-26, p=0.0313.

**Supplementary Fig. 5 | Confirmation of viral transfection with AAV2-CaMKII-NPTX2-SEP and AAV2-CaMKII-dnNPTX2-EGFP. A**, AAV-CaMKII-NPTX2-SEP was injected into the visual cortex of PV-Cre;Ai14(tdTomato) newborn mice (p0-2). After 3-4 weeks, the effect of overexpressed NPTX-SEP on Pyr→PV-IN connectivity and ODP were assessed. Expression of NPXT2-SEP in proximity of PV somas was confirmed by confocal microscopy. **B-C**, AAV-CaMKII-NPTX2-EGFP was injected into the same line of adult mice (p80-114). After 3-4 weeks, the effect of decreased expression of NPTX-SEP on Pyr→PV-IN connectivity and ODP) was assessed. The expression of NPTX2-EGFP was detected on the slices (**B**) and through the cranieal wiondow (**C**).
